# Encoding surprise by retinal ganglion cells

**DOI:** 10.1101/2022.10.15.512347

**Authors:** Danica Despotović, Corentin Joffrois, Olivier Marre, Matthew Chalk

## Abstract

The efficient coding hypothesis posits that early sensory neurons transmit maximal information about sensory stimuli, given internal constraints. A central prediction of this theory is that neurons should preferentially encode stimuli that are most surprising. Previous studies suggest this may be the case in early visual areas, where many neurons respond strongly to rare or surprising stimuli. For example, previous research showed that when presented with a rhythmic sequence of full-field flashes, many retinal ganglion cells (RGCs) respond strongly at the instance the flash sequence stops, and when another flash would be expected. This phenomenon is called the ‘omitted stimulus response’. However, it is not known whether the responses of these cells varies in a graded way depending on the level of stimulus surprise. To investigate this, we presented retinal neurons with extended sequences of stochastic flashes. With this stimulus, the surprise associated with a particular flash/silence, could be quantified analytically, and varied in a graded manner depending on the previous sequences of flashes and silences. Interestingly, we found that RGC responses could be well explained by a simple normative model, which described how they optimally combined their prior expectations and recent stimulus history, so as to encode surprise. Further, much of the diversity in RGC responses could be explained by the model, due to the different prior expectations that different neurons had about the stimulus statistics. These results suggest that even as early as the retina many cells encode surprise, relative to their own, internally generated expectations.

## Introduction

Visual scenes are highly correlated, both in space and time. It has been hypothesized that neurons in early sensory areas have evolved to exploit this structure, by only encoding ‘surprising’ sensory signals, that cannot be predicted based on their spatio-temporal context. This efficient coding theory can account for many qualitative aspects of neural responses in early sensory areas, such as the stimulus selectivity of neurons in the retina [Karklin and Simoncelli, 2011, Doi et al., 2012, Soto et al., 2020], as well as primary visual [Rao and Ballard, 1999, Olshausen and Field, 1996, Van Hateren and van der Schaaf, 1998] and auditory [Lewicki, 2002, Smith and Lewicki, 2006] cortices.

A central prediction of the efficient coding theory is that neurons should best encode stimuli that are surprising, given the recent stimulus history. There appears to be some evidence for this in early visual and auditory areas, where neurons have been found that respond most strongly to rare or surprising stimuli [Ulanovsky et al., 2003, Gill et al., 2008]. In the retina, previous studies found that when a sequence of full-field light flashes are presented, many neurons respond most strongly at the moment the sequence of flashes is stopped, and where another flash would be expected. This phenomenon was labelled the ‘omitted stimulus response’ (OSR), since it can be considered to be a response to the stimulus that was unexpectedly omitted [Schwartz et al., 2007].

However, if neurons in the retina really do encode surprise, then their responses should vary in a graded way as one varies the level of stimulus surprise. Unfortunately previous studies [Schwartz et al., 2007, Schwartz and Berry 2nd, 2008, Werner et al., 2008] did not test for this, since there were typically only two alternatives: either the stimulus was surprising (e.g the sequence of flashes ends) or it was unsurprising (e.g. the sequence of flashes continues). As a result, it is hard to conclude from these studies whether neurons in the retina encode surprise.

To address this question, we presented retinal ganglion cells (RGCs) with extended sequences of stochastically occurring full-field flashes. With this stimulus, the degree of ‘surprise’ for each flash (or period of silence between flashes) could be quantified mathematically, and was observed to vary in a graded manner depending on the previous sequence of flashes and silences. We could thus test how RGC responses varied with the level of surprise. Interestingly, we found that the responses of RGCs to these stimulus sequences could be well explained by a simple normative model, which described how neurons optimally combined their prior expectations about the stimulus with the recent stimulus history to encode surprise. Further, we found that much of the diversity in the responses of different recorded RGCs could be explained by this model, due to the different levels of ‘confidence’ that different neurons had in their prior expectations. Our study provides support for the predictive coding model of retinal coding, while shedding light on the different prior expectations that different RGCs have about environment. More generally, it shows that, already at the stage of the retina, many ganglion cells do not encode the physical stimulus itself, but how unexpected this stimulus is, with different prior expectations for different cells.

## Results

### RGC responses to flash sequences

We used a multi-electrode array to record retinal ganglion cells (RGCs) of an axolotl. We presented a visual stimulus, consisting of random sequences of full-field dark flashes, interleaved with periods of silence (Fig. 1A; see Methods: Stimulus statistics for details). Recorded neural activity was sorted into single unit responses using SpyKing Circus [Yger et al., 2016].

**Figure 1:**
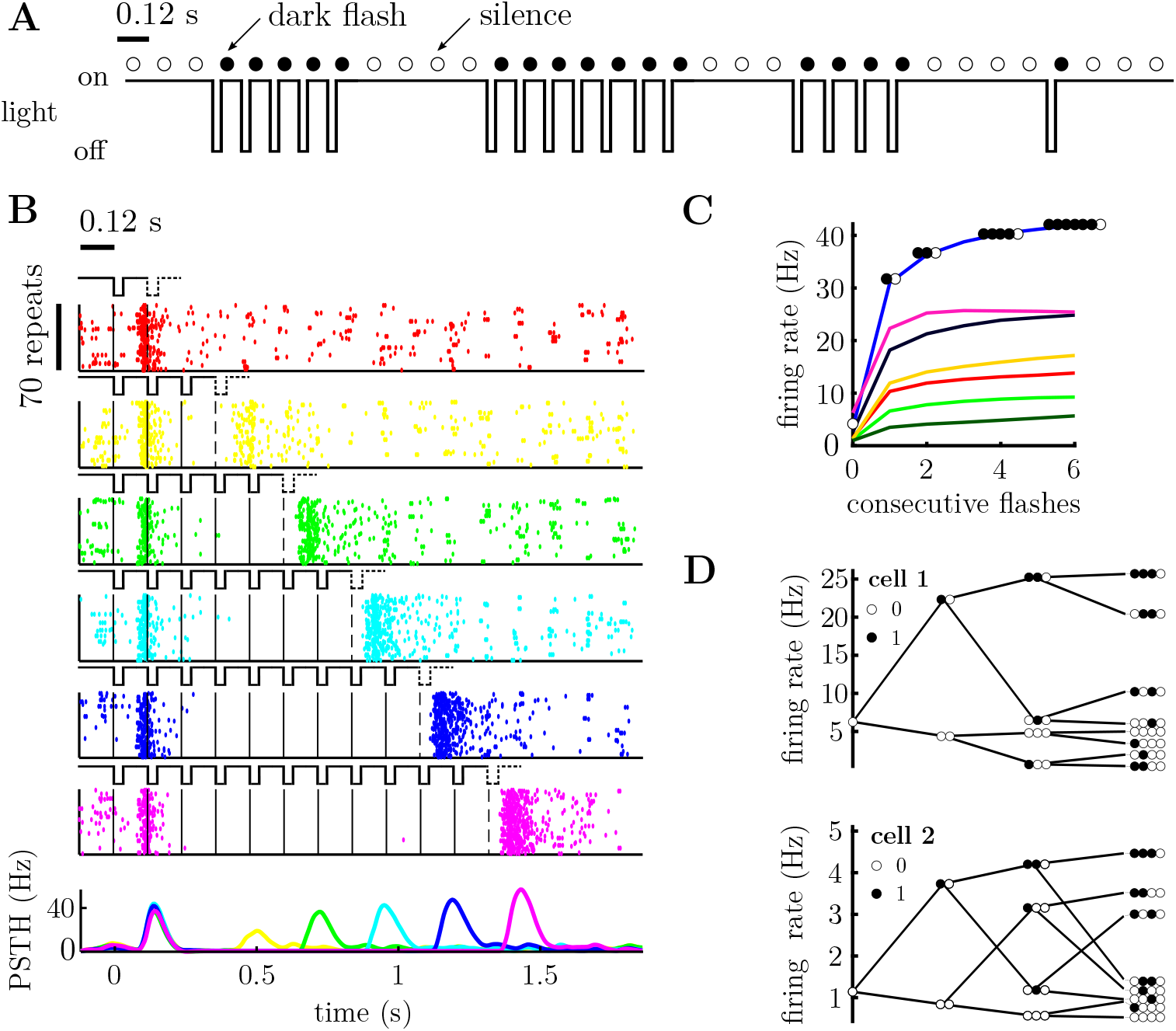
Retinal ganglion cells’ responses to a sequences of dark flashes and silences. **A**. Stimulus excerpt, showing periodic sequences of dark flashes. Each flash lasts 40 ms with 80 ms between. The 120 ms bin containing a single dark flash is marked with a filled circle. A bin without a flash (called a ‘silence’) is marked by an open circle. **B**. Raster plot for one cell. A solid vertical line marks the occurrence of each flash; a dashed line indicates an ‘omitted’ flash, following a sequence of flashes. Each raster plot shows the cell’s response to a different number of consecutive flashes (ranging from 1 to 11), with 70 repeats shown in each row of the raster. The bottom row shows the peri-stimulus time histogram (PSTH) for different numbers of consecutive flashes (colors denote the number of flashes). There is an increase in firing rate after the missing flash, called the omitted stimulus response (OSR). The OSR magnitude increases with the number of flashes. **C**. OSR for 7 cells, following a varying number of consecutive flashes (filled circles) followed by silence (open circles). **D**. Tree-plot, showing the mean response of two representative cells to different sequences of flashes (filled circles) and silences (empty circles). Cell 1 is the cell plotted in pink in panel C. Each column of the tree-plot shows the average response of the neuron to all stimulus sequences of a given length that end with silence (tree-plots corresponding to sequences ending with a flash are shown in Supp. Fig. 1). Moving right-ward the tree-plot branches out to include the effect of stimuli presented further in the past. The top branch of the tree-plot shows the cells’ responses to a series of consecutive flashes followed by silence, as in panel C. Other branches show the cells’ responses to all the different possible flash sequences of a given length.

We were interested in neurons that exhibited an ‘omitted stimulus response’ (OSR), where they responded to the absence of a flash, following several flashes presented in a row [Schwartz et al., 2007]. We thus selected 48 out of 114 single unit responses for further analysis, that showed (i) high quality recording (quantified by low number (*<*1%) of refractory period violations, where refractory period is 2 ms), and (ii) the presence of an OSR (quantified as a peak around 120 ms after the omitted flash).

Fig. 1B shows the example responses of one of these cells to a varying number of flashes presented in a row. As can be seen, this cell responded strongly to the first flash in a sequence, and shortly after the sequence had ended (i.e. the OSR). The size of the OSR increased monotonically with the number of flashes presented in a row.

For our analysis, we converted the stimulus to a binary variable, which was set to 1 or 0 depending on whether there was a dark flash (stim. = 1) or a period of silence (stim. = 0) within a given 120ms window. Neural responses were taken to be the number of spikes that occurred within each 120ms window.

To see how the OSR varied with the number of consecutive flashes, we computed the average response of each neuron, given a ‘stimulus history’ consisting of a varying number of consecutive flashes followed by silence (Figure 1C). The OSR increased monotonically with the number of flashes for all cells. However, we observed differences in the rate of increase as well as the maximum firing rate for different cells (Fig. 1C).

Finally, to see how neural responses depended on all possible stimulus sequences (and not the number of consecutive flashes), we constructed ‘tree-plots’ (Fig. 1D), showing each neuron’s average response to all possible sequences of flashes and silences of a given length. The top branch of this tree plot corresponds to the OSR, shown in Fig. 1D. However, many cells that showed a qualitatively similar OSR (i.e. that increased with the number of consecutive flashes) exhibited very different tree-plots, identifying clear differences in how they responded to different patterns of flashes and silences (e.g. Fig. 1D).

### Modeling ‘surprise encoding’ by RGCs

We asked whether RGC responses were consistent with them encoding surprise. To test this, we constructed a simple model of how RGCs could combine their internal stimulus expectations with their recent stimulus history to compute surprise (Fig. 2A). Following [Shannon, 1948, MacKay et al., 2003], we defined surprise at time *t, s*_*t*_, as the negative log probability of a stimulus, *x*_*t*_, given the recent stimulus history, *x*_*<t*_, and the neuron’s internal model of the stimulus statistics (parameterised by *θ*):

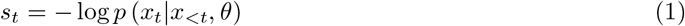

The mean firing rate was then obtained by applying a simple non-linear mapping:

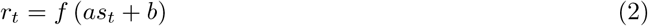

where *a* and *b* are free parameters and *f* (·) was assumed to be a softplus (log(1 + *e*^*x*^)) non-linear function to prevent firing rates being negative.

**Figure 2:**
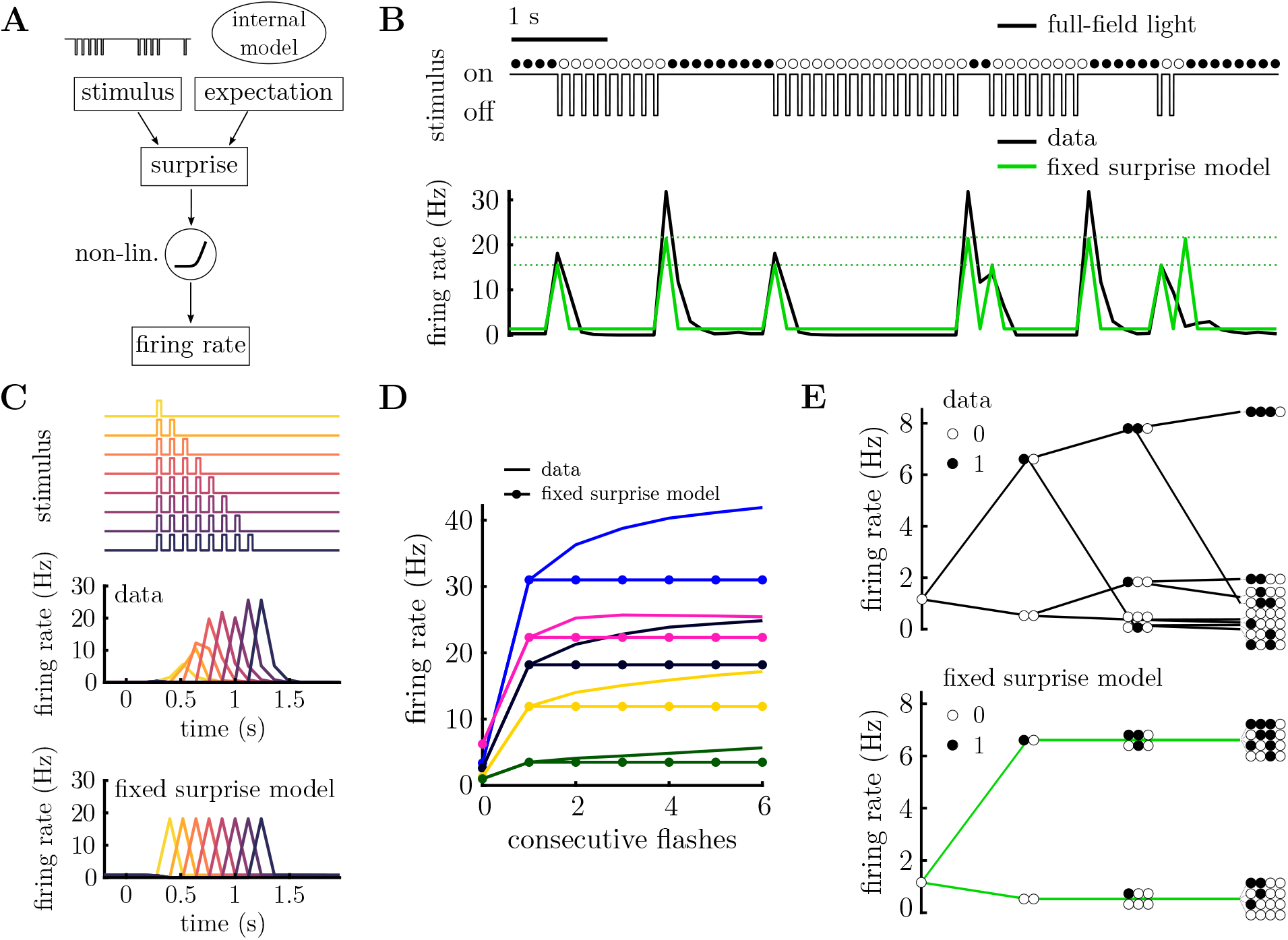
Fixed surprise model. **A**. Schematic of modeling framework. The stimulus is compared to the neuron’s expectation, which depends on their internal model, to compute surprise. The encoded surprise is then transformed via a static non-linearity to obtain the neuron’s firing rate. **B**. Stimulus excerpt (above) and recorded PSTH (below, black), and prediction of the fixed surprise model (below, green). The fixed surprise model has limited flexibility, only permitting four possible firing rates (indicated with dashed lines). **C**. Response to flash sequences of varying length (top). PSTH for a single neuron (middle) and model prediction (bottom) to the stimulus sequences shown above. Each colour corresponds to a different length of flash sequence. The fixed surprise model predicts the OSR magnitude to be independent of the number of flashes. **D**. OSR for 5 cells (solid lines) after a varying number of consecutive flashes. The fixed surprise model (lines with filled circles) cannot account for the increase in the OSR with increasing number of flashes. **E**. Tree-plot for a single cell (above) and fixed model prediction (below). The fixed surprise model can capture the mean response for stimulus sequences of up to length 2, but not beyond. Tree-plots corresponding to sequences ending with a flash are shown in Supp. Fig. 2.

The computed ‘surprise’ for each cell thus depends on their expectations or ‘internal model’ of the stimulus statistics (parameterized by *θ*). We first assumed the simplest possible internal model: a ‘Markov model’, in which the probability of observing a flash, *x*_*t*_ = 1, only depends on whether there was a flash or not in the previous time bin (*x*_*t*_ = 0*/*1). This binary Markov model has two free parameters: the probability of a flash occurring if there was/wasn’t a flash in the previous time-step (*θ*_0_ = *p* (*x*_*t*_ = 1|*x*_*t−*1_ = 0), and *θ*_1_ = *p* (*x*_*t*_ = 1|*x*_*t−*1_ = 1)). The parameters of the response function (*a* and *b*) and internal model (*θ*) were fitted for each neuron using maximum likelihood, assuming that the responses were generated by a Poisson distribution with mean *r*_*t*_ (see Methods: Neural model).

Fig. 2B shows the average firing rate of a single neuron (black) for a given stimulus sequence (above) (see Methods: Data analysis). This model accounted for the most prominent feature of the neuron’s responses: that it responded strongly to the first flash in a sequence, and the first silence in a sequence (i.e. the OSR). However, the model was unable to replicate the dependence of the OSR on the number of flashes presented in a row, observed for this (Fig. 2C) and many other cells (Fig. 2D). This was because, by design, with a Markov model the computed surprise, only depends on the stimulus in the previous time-bin, and thus the predicted response is also independent of the stimulus history, beyond one time-bin (Fig. 2E).

### Adaptive surprise model

To account for the observed variations in the OSR with number of consecutive flashes, we next considered a more complex ‘dynamic belief’ internal model. Here, we assume that the transition probabilities (*θ*_*i*_ ≡ *p* (*x*_*t*_ = 1|*x*_*t−*1_ = *i*)), are not known *a priori* by each neuron, but must be inferred. We assume neurons combine their prior expectations (*p* (*θ*_*i*_)) with the recent stimulus history (*p* (*x*_*t*_, *x*_*t−*1_, … |*θ*)) using Bayes’ law: *p* (*θ*|*x*_*t*_, *x*_*t−*1_) ∝ *p* (*x*_*t*_, *x*_*t−*1_, … |*θ*_*i*_) *p* (*θ*_*i*_). We assumed a beta-distribution for the prior over *θ*_*i*_, with parameters *α*_*i*_ and *β*_*i*_. This results in a simple expression for the inferred probability of observing *x*_*t*_ = 1, given *x*_*t−*1_ = *i*:

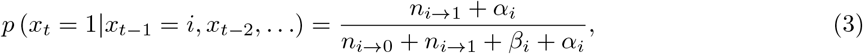

where *n*_*i*→*j*_ is the number of occurrences of the transition *i* → *j* in the sequence {*x*_1_, *x*_2_, …, *x*_*t*_}, and *α*_*i*_ and *β*_*i*_ are parameters of the prior. We assume that the parameters of the prior (*α*_*i*_, *β*_*i*_) are different for each neuron. Note that, in the limit where the prior is very strong (i.e. *n*_*i*→*j*_ ≪ *α*_*i*_ and *n*_*i*→*j*_ ≫ *β*_*i*_), this model becomes identical to the ‘fixed-belief’ model described in the previous section, where the transition probabilities for each neuron are stimulus-independent.

If neurons had ‘infinite’ memory then, given a sufficiently long stimulus sequence, their prior expectations would have no effect. Instead, we assume a more biologically plausible model where neuron’s have a finite memory, and *n*_*j*→*i*_ are estimated using a leaky integration of past observations (see Methods: Adaptive surprise model). This requires one additional parameter (the time-scale of integration i.e. the leak parameter), which we kept fixed for all neurons. In Supplementary section Dynamic surprise model we show how qualitatively similar results can be obtained by assuming neurons perform Bayesian inference, given a model where the transition probabilities have a small probability of changing on each time-step. However, optimal Bayesian inference in this setting required complex numerical integration, and it is thus hard to see how it could be implemented feasibly by individual neurons. As a result, we focus on the simpler ‘leaky integration’ model for the rest of the paper.

We fitted the 4 parameters of the prior (plus the bias and gain of the LN model) for each neuron, using maximum likelihood, assuming Poisson noise [Pillow et al., 2005]. Fig. 3A shows the predicted firing rate for one neuron (blue) to a short stimulus sequence (above). The ‘adaptive surprise’ model was able to capture aspects of the neuron’s response that could not be accounted for by the previous ‘fixed surprise model’. For example, it could capture how the size of the OSR increased with the number of flashes presented in a row (Fig. 3B-C). Further, it captured individual differences in the OSR decay for different neurons (compare cells 1-3 in Fig. 3B). Overall, the correlation between the estimated firing rates and the model prediction was significantly higher for the adaptive, compared to the fixed, surprise model (Fig. 3D).

**Figure 3:**
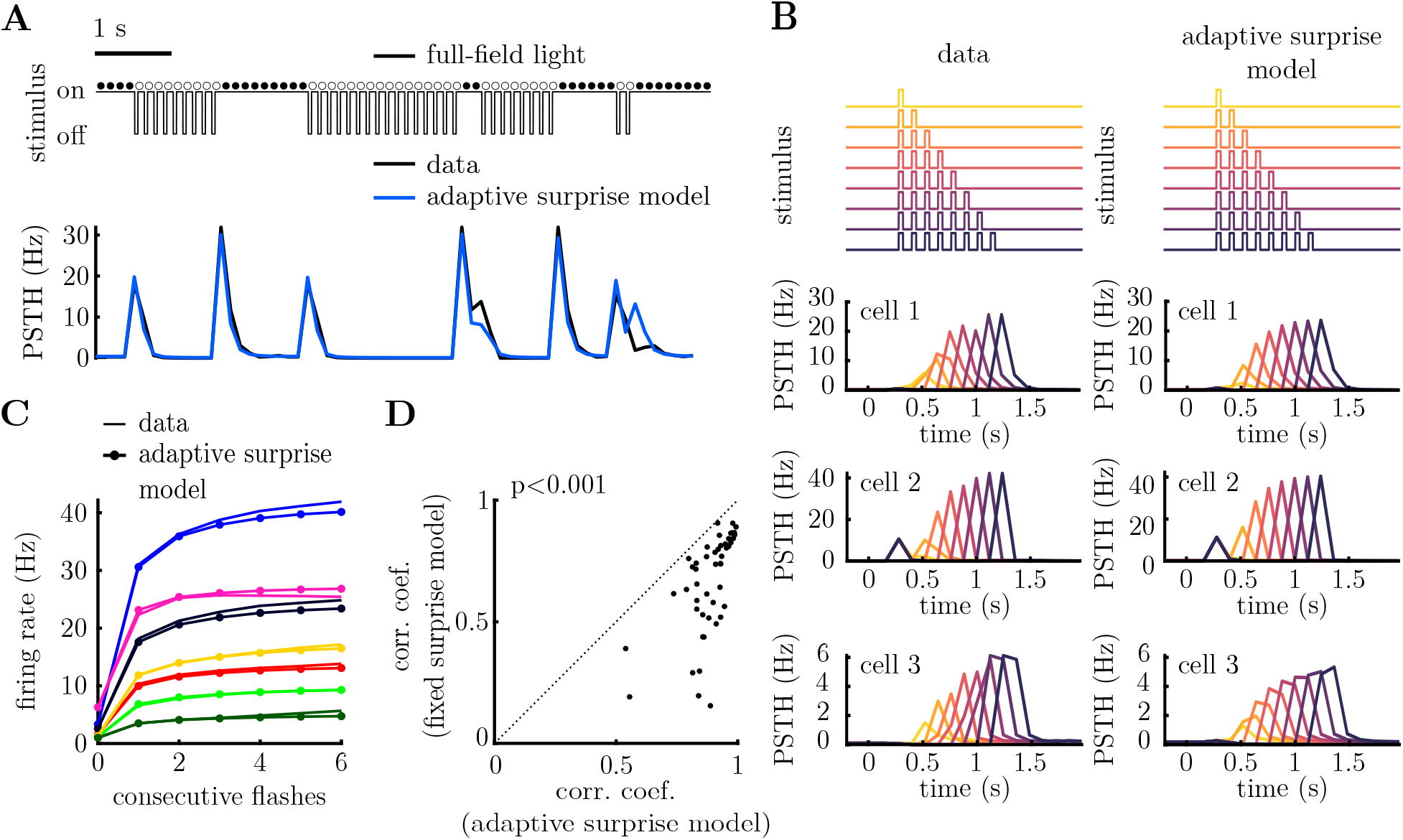
Adaptive surprise model. **A**. Stimulus excerpt (above) and recorded PSTH (below, black), alongside prediction of the adaptive surprise model (below, blue). **B**. Neural responses to varying number of consecutive flashes (above). Recorded PSTH of three neurons is shown to the left, while model predictions are shown to the right. Each colour corresponds to a different number of consecutive flashes. The adaptive surprise model captures variations in both the magnitude and width of the OSR. **C**. Increase in the OSR with the number of consecutive flashes for seven cells (each cell plotted with a different colour). The data (solid line) is plotted alongside the predictions of the adaptive surprise model (solid lines with circles). **D**. Pearson correlation coefficients between each cell’s PSTH and the model predictions, for the fixed surprise model (y-axis) versus the adaptive surprise model (x-axis). The adaptive surprise model significantly outperforms the fixed surprise model (p = 1 · 10^*−*9^, Wilcoxon signed-rank test).

To further investigate how the adaptive surprise model could account for the diverse responses of different cells, we plotted tree-plots showing the average firing rates predicted by the model for stimulus sequences of different lengths (Fig. 4A). The adaptive belief model captured much of the structure in the neural responses to stimulus sequences of varying length, as well as the diversity across different cells. This was supported by plotting the correlation coefficient between the model predictions for each node of the tree and the data, which decayed slowly with the tree depth (Fig. 4B), compared to the fixed belief model which reduced dramatically for tree depth greater than 2.

**Figure 4:**
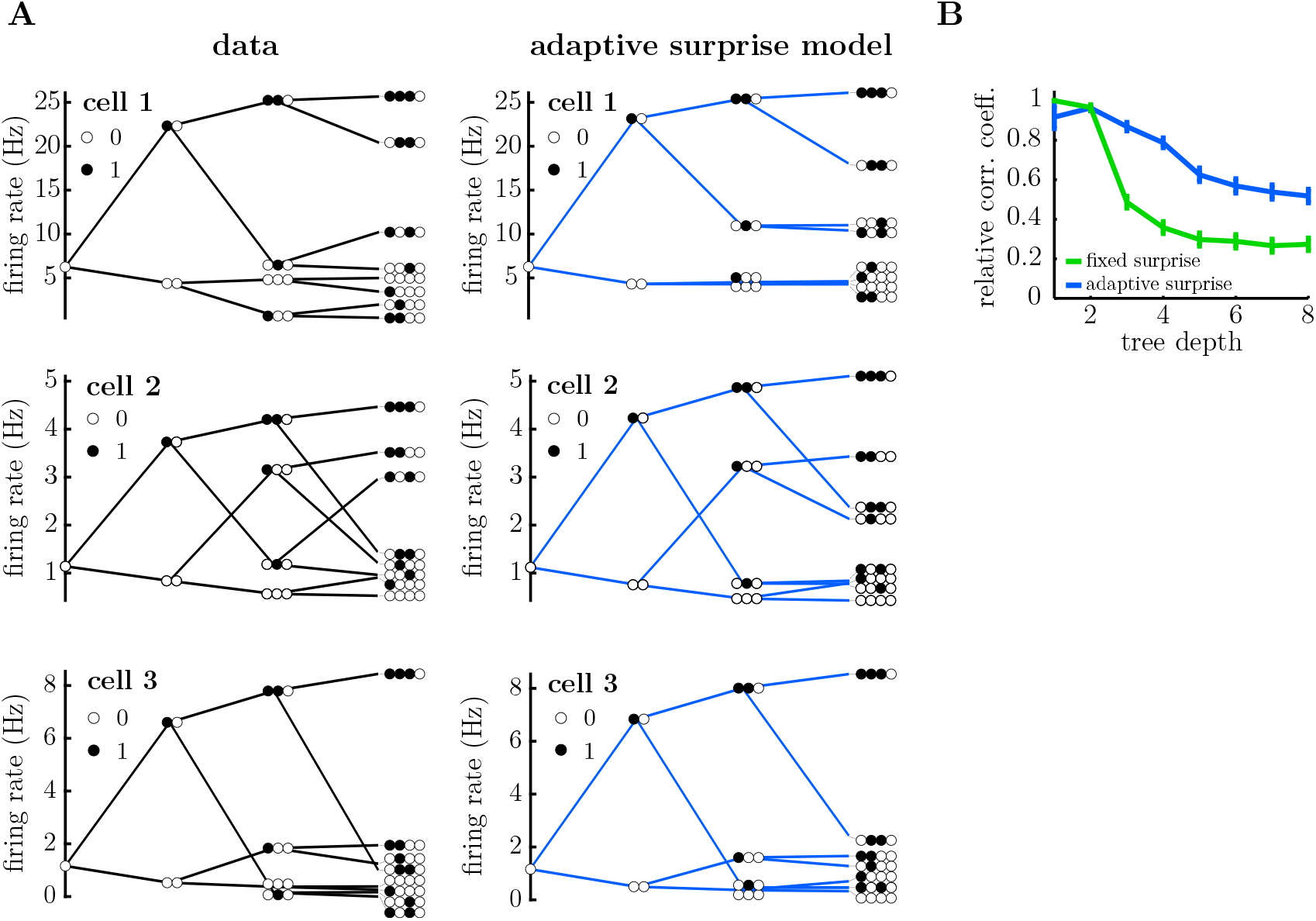
Neural responses to different stimulus sequences, for the adaptive surprise model. Tree-plot, showing the mean response of three representative cells to all possible sequences of flashes (filled circles) and silences (empty circles) of a given length. As we move rightward, the tree branches to show responses to take into account stimuli presented further in the past. The data is shown on the left and the model predictions on the right. The adaptive model is a able to reproduce qualitative aspects of each tree-plot, beyond the top branch (which shows how the OSR magnitude varies with the number of consecutive flashes). Tree-plots corresponding to sequences ending with a flash are shown in Supp. Fig. 3. **B**. Correlation coefficient between the tree-plot obtained with the adaptive surprise model and the data, computed separately for each tree-depth (i.e. stimulus sequence length). The adaptive model is significantly better at capturing the shape of the tree for stimulus sequences longer than 2, compared to the fixed surprise model.

To further test our adaptive belief model, we compared it to a more complex fixed belief model, with a comparable number of free parameters. To do this, we implemented a ‘Markov-2 model’ in which the probability of observing a flash is depends on the observed stimulus in the previous two time-bins. This model’s prior has 4 parameters, (*θ*_*ij*_ = *p*(*x*_*t*+1_ = 1|*x*_*t*_ = *i, x*_*t−*1_ = *j*)), which is the same as the adaptive surprise model (aside from the leak parameter, which we kept the same for all cells). The behaviour of this model is shown in Fig. 5. While the Markov-2 model outperformed the fixed surprise model model described earlier, it could not account for increases in the OSR that occurred for sequences of more than 2 consecutive flashes (Fig. 5B-C), or any structure in the tree plots at a depth greater than 2 (Fig. 5D, emphasized with a dashed ellipse). Finally, the correlation coefficient between predicted and observed firing rates was significantly worse for the Markov-2 model than the adaptive surprise model (Figure. 5E) despite them having the same number of free parameters (*p* = 2 · 10^*−*8^, Wilcoxon signed-rank test).

**Figure 5:**
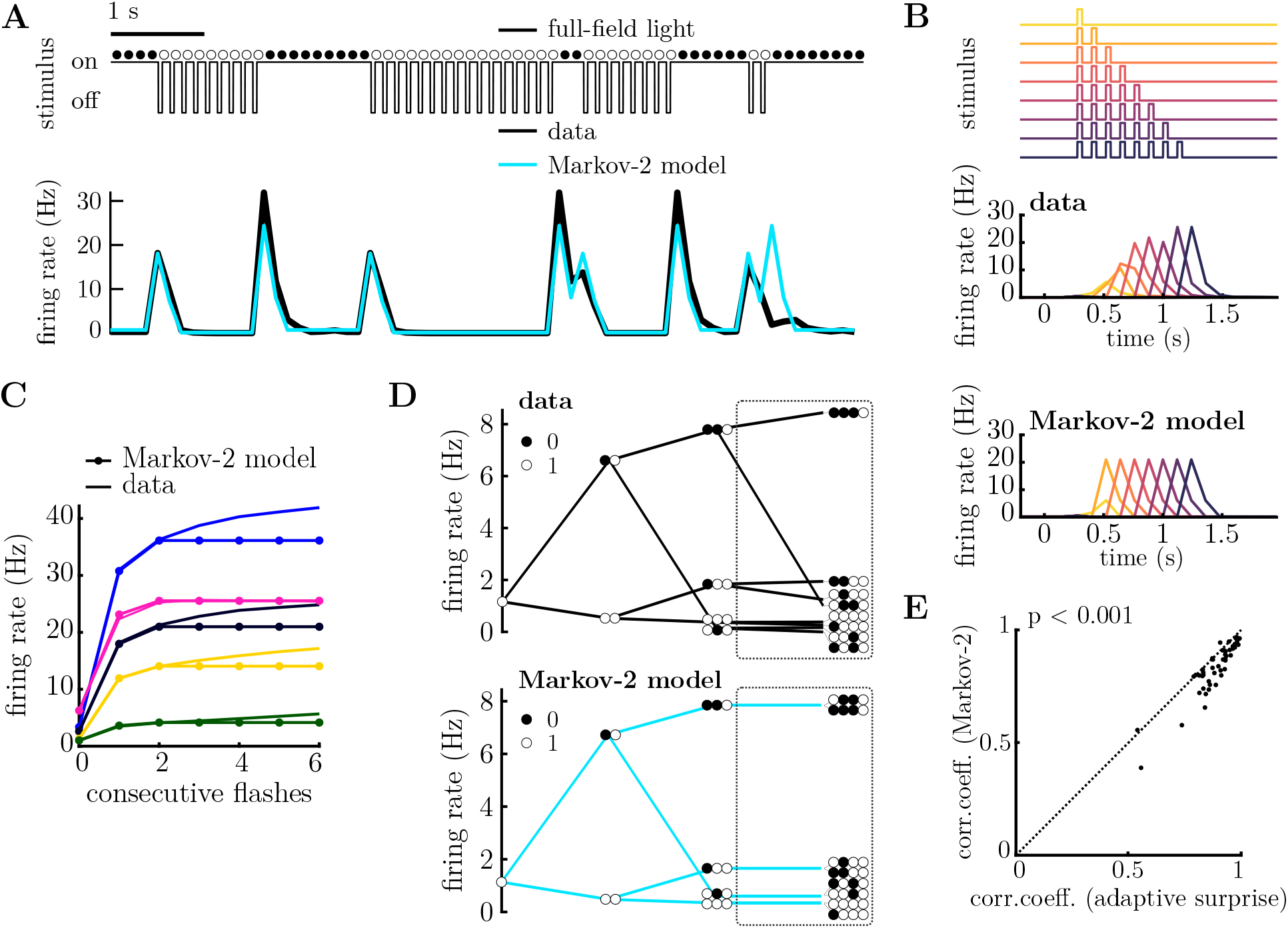
Fixed surprise model with longer past (Markov-2 model). **A**. Stimulus excerpt (above) and recorded PSTH (below, black), and firing rate predicted by the Markov-2 model (below, blue). Response to varying number of consecutive flashes (top). PSTH for a single neuron (middle) and model prediction (below) to the stimulus sequences shown above. Each colour corresponds to a different length of flash sequence. The Markov-2 surprise model predicts the OSR magnitude to be dependent on the previous two state only. **C**. Average responses of 5 cells (solid lines) to flash sequences of varying lengths. The Markov-2 surprise model (lines with filled circles) cannot account for the increase in the OSR beyond 2 consecutive flashes. **D**. Tree-plot for a single cell (above) and fixed model prediction (bottom). The Markov-2 surprise model cannot capture the response for stimulus sequences greater than length 2 (highlighted with dashed circle). Tree-plots corresponding to sequences ending with a flash are shown in Supp. Fig. 4. **E**. The correlation coefficient between the adaptive surprise model and recorded PSTH for each cell is significantly better than for the Markov-2 model (p = 2 · 10^*−*8^, Wilcoxon signed-rank test).

### Differences in the internal expectations for individual cells

We were interested to see how the inferred expectations (the ‘prior’) varied for each cell. Recall that we assumed a beta-prior over the transition probabilities *θ*_*i*_ = *p* (*x*_*t*_ = 1|*x*_*t−*1_ = *i*), with parameters *α*_*i*_ and *β*_*i*_. The mean of this prior is determined by the ratio of these two parameters, *α*_*i*_*/β*_*i*_, while its width (i.e. the level of prior uncertainty) is determined by their sum, *α*_*i*_ + *β*_*i*_. Figure 6A shows how the parameters of the prior varied for different cells. Interestingly, we found that while for different cells there was a large variation in the sum, *α*_*i*_ + *β*_*i*_, the ratio, *α*_*i*_*/β*_*i*_, was relatively constant. Thus, while the width of the prior, which determines how much weight is accorded to prior expectations versus new observations, varied greatly across cells, the prior mean was roughly constant.

**Figure 6:**
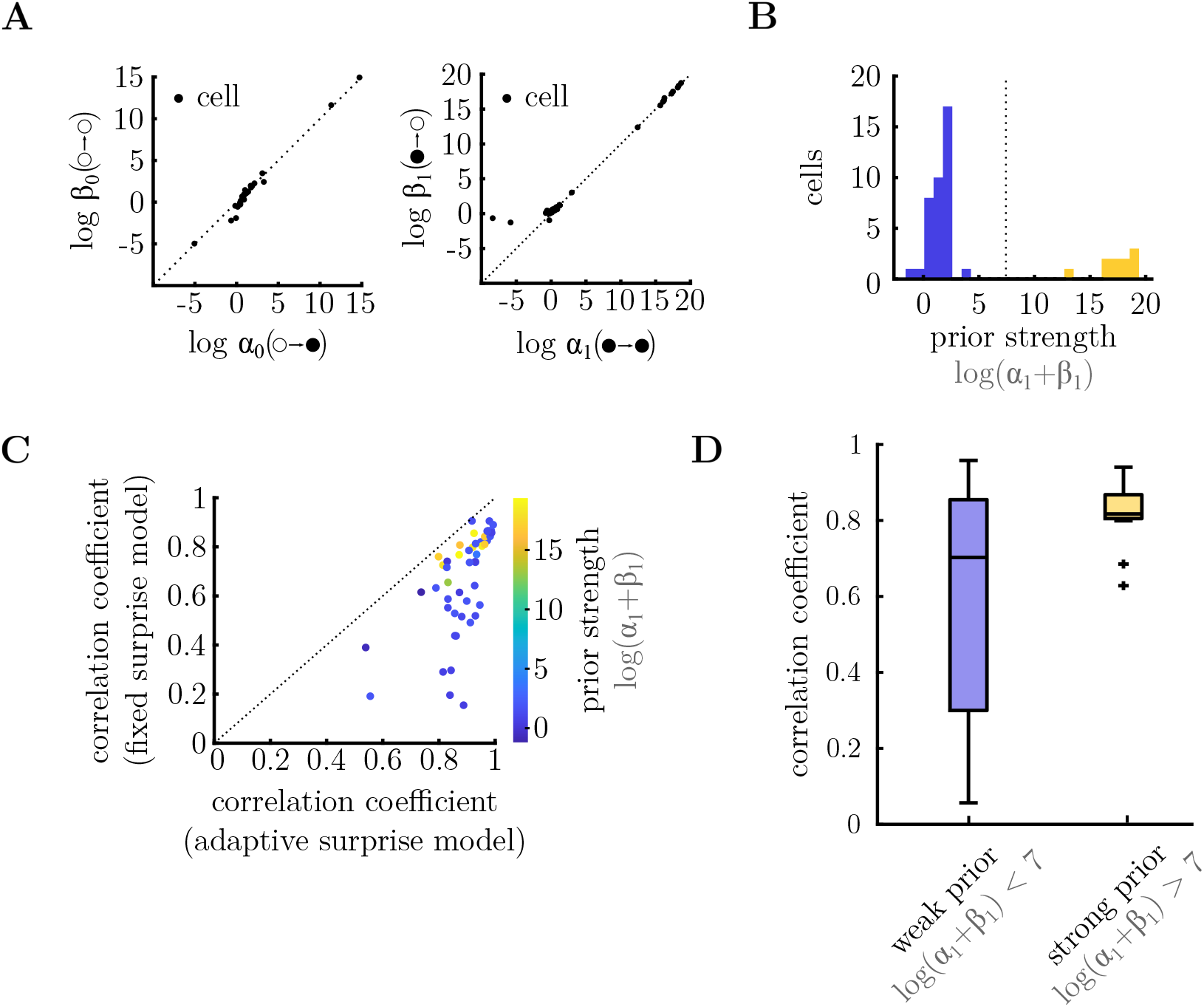
Parameters of internal model. **A**. Parameters of the inferred prior (*α*_*i*_ and *β*_*i*_) for each cell. These parameters determine each cell’s prior expectation for the different transitions from silence (left) or flash (right). In both cases, the ratio between these parameters, *α*_*i*_*/β*_*i*_ (which determine the mean of the prior) is close to unity for all the cells, while their sum, *α*_*i*_ + *β*_*i*_ (which determines the strength of the prior) varies across different cells. **B**. Histogram of log (*α*_1_ + *β*_1_) for different cells. The population could be split into two groups: cells with low confidence in the prior (i.e. small *α*_*i*_ + *β*_*i*_; blue) and cells with high confidence in the prior (i.e. large *α*_*i*_ + *β*_*i*_; yellow). **C**. Correlation coefficient between the responses predicted by the adaptive surprise model, versus the fixed surprise model. Each circle is colour coded according to the parameters of the inferred prior for that cell log(*α*_1_ + *β*_1_). Cells that had a strong prior (yellow) tended to be better fit by the fixed surprise model, relative to the adaptive surprise model. **D**. For each cell we computed the correlation coefficient between the average neural responses to stimulus sequences of length 2, versus average responses that take into account stimulus sequences of length 10. The responses of cells with a strong prior (right, yellow), but not a weak prior (left, blue), could be reasonably well predicted just by looking at the most recent stimulus transition.

Focusing on the prior parameters, *α*_1_ and *β*_1_, which determines neural responses to ‘flash→flash’ and ‘flash→no-flash’ transitions (i.e. the OSR), we observed two clusters of cells (Fig. 6A, right panel), with different levels of prior uncertainty (determined by the sum, *α*_1_ + *β*_1_; Fig. 6B). We asked what effect this would have on these cells responses. We reasoned that cells with a strong prior (i.e. large *α*_*i*_ + *β*_*i*_) would not adapt their posterior belief much depending on recent observations, and hence their responses would be well predicted by the fixed surprise model. In contrast, cells with a weak prior (i.e. small *α*_*i*_ + *β*_*i*_) would be strongly influenced by recent observations, and thus their responses would be poorly predicted by the fixed-surprise model. This turned out to be the case. Fig. 6C shows the correlation coefficient between the prediction of the fixed surprise model and recorded responses, versus the adaptive surprise model. There was a trend for cells with a strong prior (high *α*_*i*_ + *β*_*i*_; colour coded in yellow) to be equally well-fit by both models, while cells with a weak prior (low *α*_*i*_ + *β*_*i*_; colour coded in blue) were better fit by the adaptive surprise model. We asked whether this same effect could be observed without reference to the model fits. To do this, we compared the average neural responses to stimulus sequences of length 2, to the average response to longer sequences, of length 10. As expected, we found that the average responses of cells with a strong prior (i.e. log (*α*_1_ + *β*_1_) *>* 7) only depended on the most recently presented stimuli (Fig. 6D, yellow). This tended not to be the case for stimuli with a weak prior (Fig. 6D, blue) (p = 0.066, Wilcoxon rank-sum test).

In Fig. 6A we observed that the prior mean, determined by *α*_*i*_*/β*_*i*_, remained near-unity across different cells. We thus, asked whether it would be possible to fit neural responses using a reduced adaptive surprise model with only two parameters (i.e. the sum, *α*_*i*_ + *β*_*i*_), and *α*_*i*_*/β*_*i*_ held fixed at unity. Fig. 7A shows that, while this reduced adaptive model performed worse than the full adaptive surprise model, this reduction in performance was small (*<* 10% reduction in correlation coefficient), despite having having only 2 free parameters per cell (compared to 4 parameters, for the full model). Notably, the reduced model was able to capture similar qualitative features of neural responses, such as how the OSR increased with the number of consecutive flashes (Fig. 7B, Supp. Fig. 5). Its performance was also significantly better than the fixed surprise model, which had the same number of free parameters per cell (p = 4 · 10^*−*5^, Wilcoxon signed-rank test).

**Figure 7:**
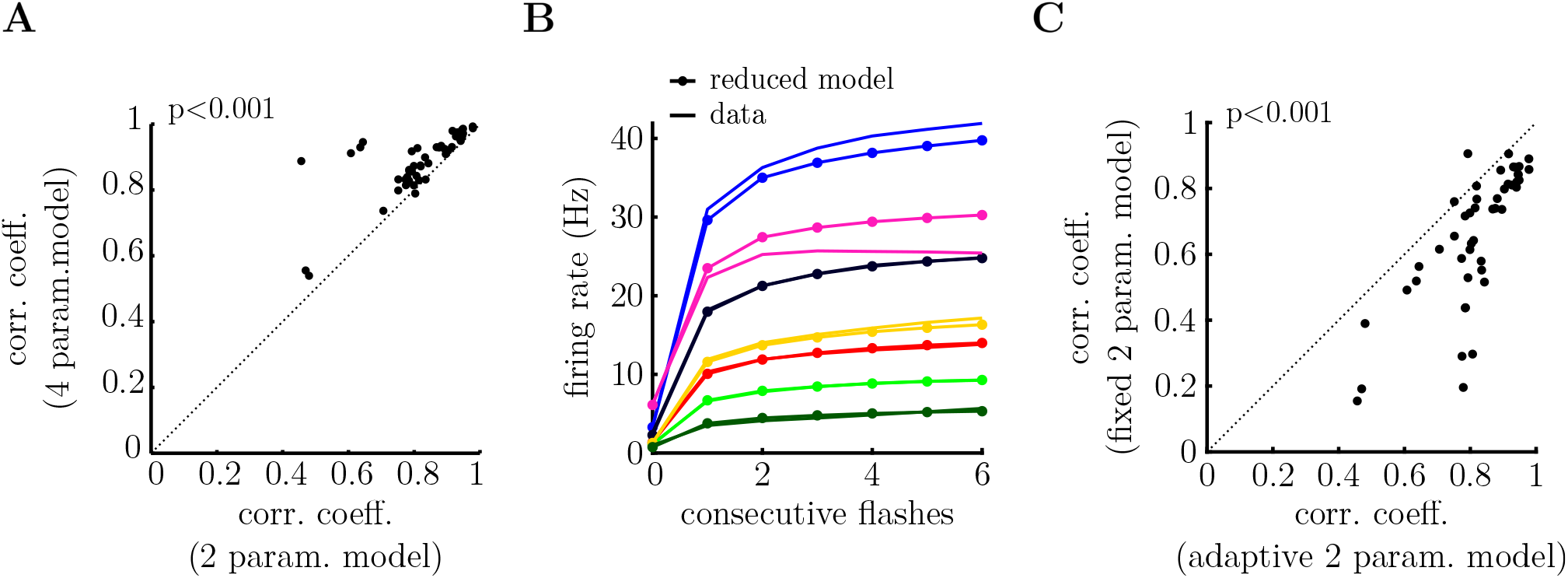
Reduced adaptive surprise model, with a fixed prior mean (i.e *α*_*i*_*/β*_*i*_ = 1). **A**. The reduced model performs almost as well as the adaptive surprise model despite having half the number of free parameters. (p = 3 · 10^*−*9^, Wilcoxon signed-rank test). **B**. Mean response of 7 cells following a variable number of flashes presented in a row. The increase in the OSR with the number of flashes is well-fitted by the reduced model. **C**. The reduced adaptive surprise model performs significantly better at fitting the recorded neural responses, despite both models having the same number of free parameters per cell. (p = 4 · 10^*−*5^, Wilcoxon signed-rank test.)

## Discussion

We observed how neural responses in the retina showed non-trivial dependencies on the precise order of flashes and silences in random stimulus sequences (Fig. 1C-D). Interestingly, RGC responses were well predicted by a simple model, which assumed that they depended on how ‘surprising’ stimuli were, relative to an internally generated expectation (Fig 3-4). Moreover, our model showed how the different ‘expectations’ of different neurons could account for the diverse way they responded to presented stimuli (Fig 6-7).

Our approach contrasts with previous ideal observer models, which assume that neurons are perfectly adapted to the ‘true’ presented stimulus statistics [Geisler, 1989, Geisler, 2003, Smeds et al., 2019, Chichilnisky and Rieke, 2005]. Instead, we found that neural responses could be well explained by assuming that each neuron has learned its own internal model of the stimulus statistics (with the parameters of the prior fitted separately for each cell). Interestingly, we found that different neurons had very similar prior expectations about which stimuli were most likely to occur (determined by the mean of the prior). What varied was the degree of confidence they had about their own prior expectations (determined by the width of the prior). Furthermore, recorded cells could be divided into two categories: those with weak confidence in their prior, and those with strong confidence in their prior expectations. In the future, it would be interesting to elucidate the reason for this split, and whether, for example, it corresponded to different types of ganglion cell identified in previous work [Baden et al., 2016].

Our modelling framework was adapted from a previous model of Meyniel et al., that sought to explain psychophysical data showing how subjects’ behaviour (such as their reaction time and accuracy) depended on the statistics of sequentially presented sensory stimuli [Meyniel et al., 2016]. Meyniel and colleagues showed how their data could by explained if subjects used a Bayesian inference model, as described here, to predict new stimuli based on what came before. Here, we extended this model to include a variable ‘prior’ distribution, whose parameters could be fit to describe the diverse responses of different ganglion cells. Nonetheless, the fact that a similar type of model can be used to describe both neural responses in the retina and subjects behaviour in different tasks is intriguing, raising the question of whether similar computations may be present ubiquitously in the brain when subjects are presented stimuli with complex temporal statistics.

Previous experimental [Schwartz and Berry 2nd, 2008, Werner et al., 2008, Deshmukh, 2015] and computational [Maheswaranathan et al., 2019, Tanaka et al., 2019, Chen et al., 2017] studies sought to understand the neural mechanisms underlying the OSR. However, there remains some controversy over which of the proposed theories could explain all of the experimentally observed features of the OSR, such as e.g. the fact that the delay before the OSR varies linearly with the time between flashes. Our work provides further constraints to distinguish between different theories, by showing how the OSR varies depending on the precise sequence of flashes and silences (Fig 1).

The stimuli in our experiment, which consisted of sequences of full-field flashes, were chosen to be sufficiently rich so as to permit many different levels of ‘surprise’, while simple enough to permit a straight-forward analysis of neural responses. Nonetheless, in the future it would be interesting to investigate neural responses to more naturalistic stimuli, which for example, varied spatially as well temporally [Keller et al., 2012]. This would allow us to investigate, for example, the degree to which neurons’ internal model is adapted to the statistics of natural scenes, as predicted by the efficient coding hypothesis [Machens et al., 2005].

## Methods

### Experimental setup

The recordings were performed in the axolotl retina, using a multi-electrode array with 252 electrodes with 60 *μ*m spacing (procedure described in detail in [Marre et al., 2012]). The experiment was performed in accordance with institutional animal care standards of Sorbonne Université. The raw signal, recorded at 20 kHz sampling rate, was high-pass filtered at 100 Hz and then sorted offline using SpyKing Circus software [Yger et al., 2016]. The stimulus consisted of full-field dark flashes. The reason for using dark flashes was the dominance of OFF type cells in axolotl. The dark flashes had a duration of 40 ms, with 80 ms period between the flashes, (∼12 Hz frequency) (as in [Schwartz et al., 2007]).

### Stimulus statistics

We generated sequences of flashes and silences (i.e. where no flash occured in a 120ms window) using a stochastic model. The number of flashes and periods of silent states presented in a row was drawn from a negative binomial distribution, with parameters *r* and *p*. In the case of flashes, we varied the first parameter, *p*, at 20 minute intervals between three different values (0.98, 0.8 and 0.01, consecutively). The second parameter, *r* was adjusted so as to maintain a constant mean, of 7 flashes presented in a row. The length of the silence sequence was drawn from a geometrical distribution with a fixed mean *p* (*p* = 9). Changing *p* alters the degree to which the distribution is clustered around the mean. However, we observed no difference in the neural responses recorded with different values of *p*. As a result we concatenated data from neural responses to all three stimulus distributions for the rest of our analysis.

### Data analysis

To generate the spike raster plots shown in Fig. 1, we aligned the spiking responses of neurons to a sequence of *n* flashes presented in a row. The peri-stimulus-time-histogram (PSTH) plotted in the bottom row of Fig. 1 was computed by averaging the spike count recording over all the stimulus repeats, and then averaging over a 5 ms time bin.

For the remainder of the analysis, we discretised the neural responses and stimulus into time bins of length 120 ms (the time between consecutive flashes). The stimulus presented in each time-bin was treated as a binary variable: ‘1’ if there was a (dark) flash, ‘0’ otherwise. The average firing rate in each bin was computed by average the spike count over all repetitions of a stimulus sequence of length *n*. Except where stated explicitly in the text, we set *n* = 8 (so there were 256 distinct stimulus sequences used to compute the average firing rate).

### Neural model

For our model, we assumed that at each time-bin, *t*, neurons fire spikes drawn from a Poisson distribution with mean, *λ*_*t*_, given by:

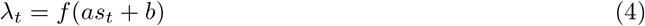

where *s*_*t*_ is the encoded surprise at time *t, f* is a non-linearity, and *a* and *b* are parameters describing the gain and bias, respectively. The non-linearity, *f* (*x*) = log(1 + *e*^*x*^) (soft-ReLU), was kept fixed for all the cells.

The surprise at time *t* is defined as:

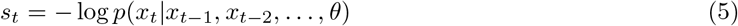

where *p* (*x*_*t*_|*x*_*t*_, *x*_*t−*2_, …, *θ*) is the probability of observing no flash or a flash at time *t* (*x*_*t*_ = 0 or 1 respectively) given the stimulus *x* at previous times, and the internal model of the cell, parameterized by *θ*.

### Internal model

The computed surprise depends on each cell’s internal model of the stimulus statistics. We first considered a binary Markov model, where the probability of observing a flash at time *t* is assumed to depend only on whether a flash was observed in the previous time bin. This model has two parameters: *θ*_0_ = *p* (*x*_*t*_ = 1|*x*_*t−*1_ = 0), and *θ*_1_ = *p* (*x*_*t*_ = 1|*x*_*t−*1_ = 1). For the Markov 2 model, we simply extend the observed history to 2 previous states, yielding a total of 4 parameters: *θ*_0_ = *p* (*x*_*t*_ = 1|*x*_*t−*1_ = 0, *x*_*t−*2_ = 0), *θ*_1_ = *p* (*x*_*t*_ = 1|*x*_*t−*1_ = 1, *x*_*t−*2_ = 0), *θ*_2_ = *p* (*x*_*t*_ = 1|*x*_*t−*1_ = 0, *x*_*t−*2_ = 1), and *θ*_3_ = *p* (*x*_*t*_ = 1|*x*_*t−*1_ = 1, *x*_*t−*2_ = 1).

### Inferring the transition probabilities

Next, we considered an ‘adaptive belief model’ where the transition probabilities, *θ*_*i*_ = *p* (*x*_*t*_|*x*_*t−*1_ = *i*), are not known in advance, but must be inferred by combining each cell’s prior belief with newly observations, using Bayes’ law, as follows:

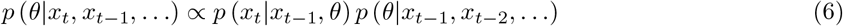

where *p* (*x*_*t*_|*x*_*t−*1_, *θ*) is the likelihood of observing *x*_*t*_ given *x*_*t−*1_, and is described by a Bernoulli distribution:

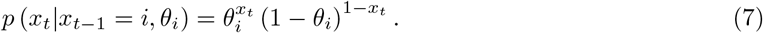

Let us assume that at time *t* − 1, the posterior distribution over *θ*_*i*_, *p* (*θ*|*x*_*t−*1_, *x*_*t−*2_, …), is described by the beta distribution with parameters 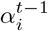 and 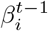:

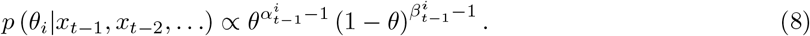

Now, at time *t*, multiplying this distribution by the likelihood according to Bayes law (Eqn 6) will result in a new beta-distribution, with parameters:

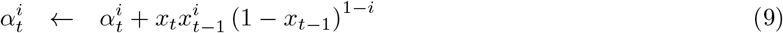

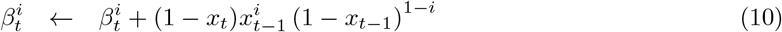

The probability of observing *x*_*t*_ = 1 given previous observations is then given by:

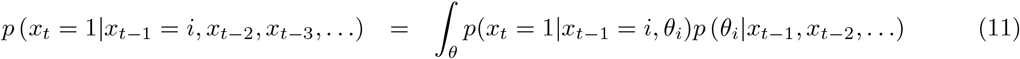

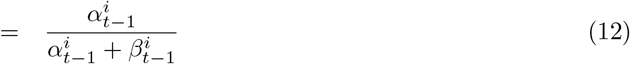

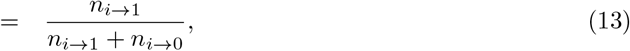

where *n*_*i*→0_ and *n*_*i*→1_ describe the number of occurrences of the transitions *i* → 0 and *i* → 1, respectively.

### Adaptive surprise model

The statistics of the external world are not static, but change in time. To take this into account, we could assume there a non-zero probability of transition matrix changing between two observations (a ‘dynamic belief model’). Performing exact Bayesian inference in this case requires expensive numerical integration, which may be difficult to perform by individual neurons in the retina. However, in [Meyniel et al., 2016] they found that such a dynamic belief model could be approximated by a ‘forgetful’ model, where recent observations are weighted more strongly than the ones in the past. In contrast to the optimal Bayesian model, their leaky integration model results in simple linear parameter updates, and could thus be easy to implement neurally.

In practice, we can implement the ‘forgetful’ model of Meyniel et al., by modifying the update rules described earlier for *α*^*i*^ and *β*^*i*^ as follows:

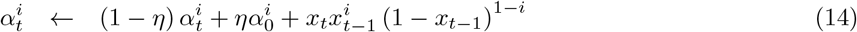

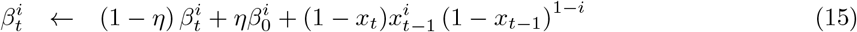

where 1 *> η >* 0 is a leak term that results in forgetting observations far in the past, while 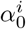 and 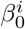 determine the steady state values of 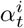 and 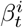 in the absence of new observations. With this update rule, the probability of observing a *x*_*t*_ = 1 given *x*_*t−*1_ = *i* is given by:

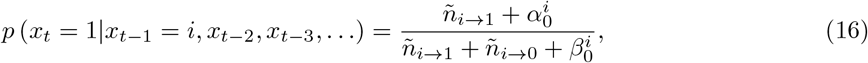

where *ñ*_*i→j*_ is the ‘effective’ number of observations of a transition *i* → *j*, after taking into account the leak, when *η >* 0:

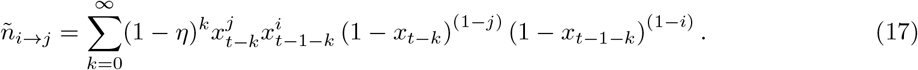

In practice we assumed that the leak, *η* was the same for all cells. We used a value of *η* = 0.2. However, similar results were obtained when we increased or decreased the leak by a small amount.

In contrast, the parameters of the prior, 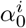 and 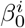, were allowed to vary for different cells. This us allowed to investigate how different cells’ ‘prior expectations’ for different transitions affected their responses. (Note that for notational simplicity we dropped the subscript ‘0’ in the main text.)

### Model fitting

We fitted the internal model parameters (see previous section), and the gain and bias of the response curves (Eqn 4) using Maximum Likelihood (ML) algorithm [Doya et al., 2007]. For this, we assumed a Poisson noise model, resulting in a log-likelihood:

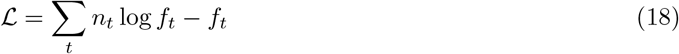

where *f*_*t*_ and *n*_*t*_ are the spike count predicted by the model and observed spike count at time *t*, respectively. All models were fitted using algorithms with multiple starting points (MultiStart in MATLAB, 50 starting points, random initial parameters).

The data analysis and model fitting were done in MATLAB R2021a. Code and data will be available upon paper acceptance.

## Acknowledgments

This work was funded by ANR JCJC grant Optimal predictive coding. The authors would like to thank Yannick Andéol for providing the axolotls. We would also like to thank Ulisse Ferrari, Francesco Trapani, Matías Goldin and Samuele Virgili for useful discussions.

