## Supplementary Information for "Encoding surprise by retinal ganglion cells"

### Complementary tree-plots

In this section, we show the tree-plots corresponding to sequences ending with a flash, corresponding to Figs 1-4.

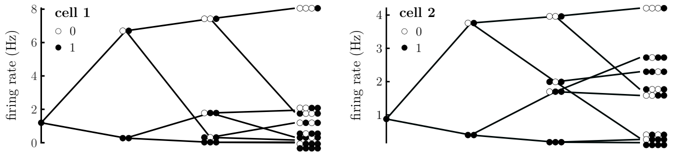


Supp. Fig. S1: Tree-plot for two representative cells, showing the mean response of two representative cells to different sequences of flashes (filled circles) and silences (empty circles). Each column of the tree-plot shows the average response of the neuron to all stimulus sequences of a given length, that end with flash.

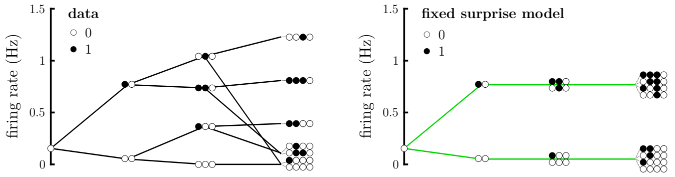


Supp. Fig. S2: Tree-plot for a single cell (left) and fixed surprise model prediction (right). Each column of the tree-plot shows the average response of the neuron to all stimulus sequences of a given length, that end with flash.

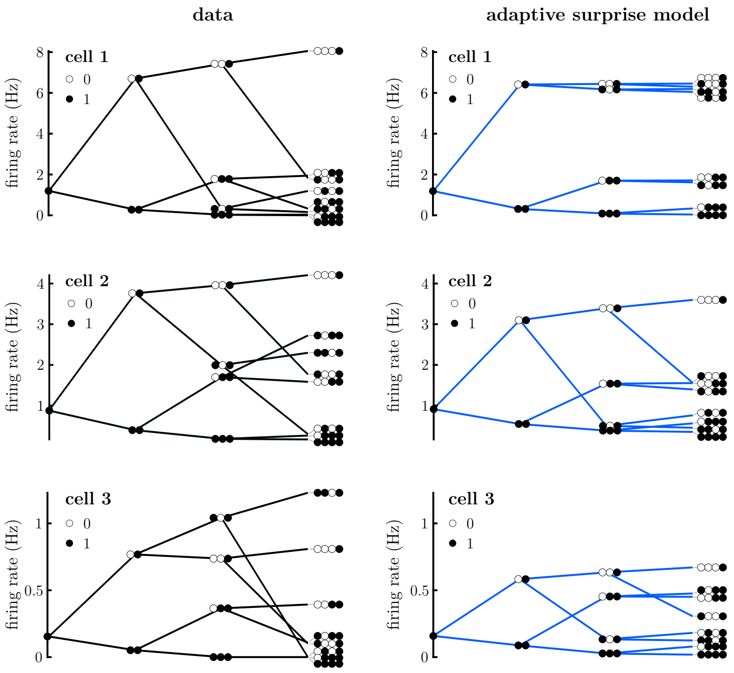


Supp. Fig. S3: Tree-plot for three representative cells, showing neuron's response (left column), and prediction by the adaptive surprise model (right column).

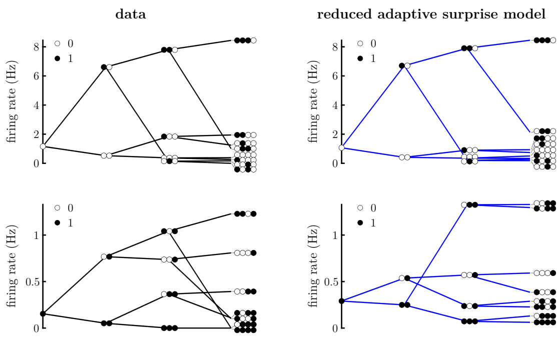


Supp. Fig. S4: Tree-plot for reduced adaptive surprise model, showing neuron's response (left column), and prediction by the adaptive surprise model (right column). The qualitative traits of the adaptive surprise model remain even if two instead of four prior parameters are used.

### Dynamic surprise model

For the dynamic surprise model, the assumption is that the transition probability of a sequence can change at each time-step with a certain non-zero probability, $p_c$. However, as discussed in the main text, this model is both biologically implausible and computationally expensive, therefore the leaky integration model was used as its approximation. First, the probability $p_c$ was learned independently for each cell. To better compare the performance of dynamic model with the adaptive model with fixed delay, the value of probability $p_c$ was set to constant for all the cells and set at median value ($p_c = 0.2$). This value was the median of $p_c$. The model was fitted using algorithms with multiple starting points (MultiStart in MATLAB, 50 starting points, random initial parameters).

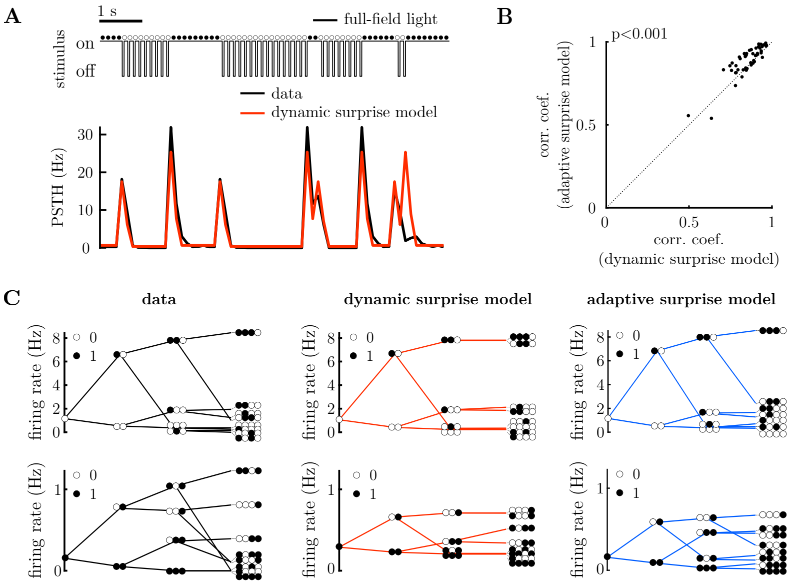


Supp. Fig. S5: **Dynamic inference model performs similar to its leaky integration approximation. A.** Stimulus excerpt (above) and recorded PSTH (below, black), and prediction of the dynamic surprise model (below, red). **B.** Pearson correlation coefficients of model fits to each cell's PSTH, for the dynamic surprise model (x-axis) versus the adaptive surprise model (y-axis). Two model perform similarly (the correlation coefficents for the respective model fits lie close to the unity line) despite the significant difference in mean ($p = 3 \cdot 10^{-5}$, Wilcoxon signed-rank test). As such, the adaptive surprise model can be treated as an approximation of the dynamic surprise model. **C.** Tree-plot, showing the mean response of two representative cells to different sequences of flashes (filled circles) and silences (empty circles). Each column of the tree-plot shows the average response of the neuron to all stimulus sequences of a given length, that end with silence (top) or flash (bottom).
